# A Message Passing Framework for Precise Cell State Identification with scClassify2

**DOI:** 10.1101/2024.06.26.600770

**Authors:** Wenze Ding, Yue Cao, Xiaohang Fu, Marni Torkel, Jean Yang

## Abstract

In single-cell analysis, the ability to accurately annotate cells is crucial for downstream exploration. To date, a wide range of approaches have been developed for cell annotation, spanning from classic statistical models to the latest large language models. However, most of the current methods focus on annotating distinct cell types and overlook the identification of sequential cell populations such as transitioning cells. Here, we propose a message-passing-neural-network-based cell annotation method, scClassify2, to specifically focus on adjacent cell state identification. By incorporating prior biological knowledge through a novel dual-layer architecture and employing ordinal regression and conditional training to differentiate adjacent cell states, scClassify2 achieves superior performance compared to other state-of-the-art methods. In addition to single-cell RNA-sequencing data, scClassify2 is generalizable to annotation from different platforms including subcellular spatial transcriptomics data. To facilitate ease of use, we provide a web server hosting over 30 human tissues.

## Introduction

Leveraging advancements in sequencing technologies, single-cell transcriptomics has revolutionised biological and medical research by offering unprecedented opportunities to observe gene expression across the entire transcriptome at cellular resolution(1–5). A fundamental task in single cell research is cell annotation, as understanding the identity of cells is key to further downstream analysis(6–8). Current cell annotation tasks are based on two main categories of approaches, supervised and unsupervised (including some semi-supervised ones)(9–16). Firstly, with an unsupervised approach, the process typically involves clustering the query data into groups based on their molecular characteristics such as gene expression. This is then followed by annotating cells from each group via marker gene analysis. The alternative approach is supervised classification, where a set of cell expression data and their corresponding known cell type labels are used as the training or reference data to build a model. The model can then be used to annotate cell types in query or unseen datasets.

Although numerous approaches for cell type annotation exist, a significant portion of these focuses on discrete and non-sequential cell subpopulations and are not designed to effectively model the relationship of the transition between cell states. In the biological system, many processes often involve cell transitions (Fig. 1a), such as human preimplantation embryo development(17) and T cell differentiation during infection(18). In contrast to non-sequential cell type annotation problems, sequential cell states are typically more challenging due to the similar nature between adjacent cell states (Fig. 1b). There is a gap of existing approaches specially designed to account for cell state transition and thus effectively discriminate adjacent cell states remains a challenge in current cell annotation(19, 20).

**Fig. 1.**
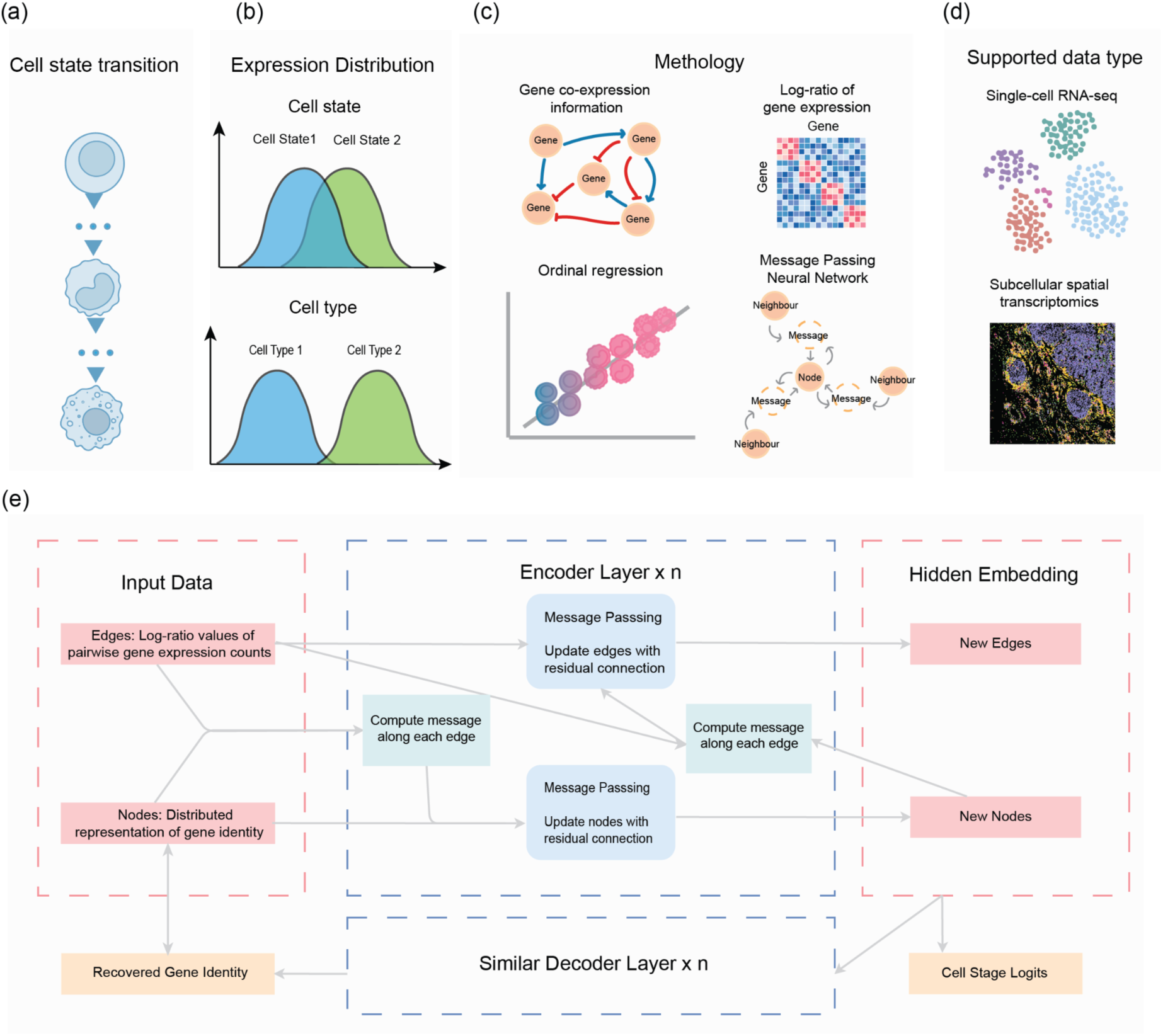
Overview of the scClassify2 framework for sequential cell state identification. (a) Linear cell state transformation is one of the most common and important phenomena in nature when external disturbance factors like signal molecules, drugs or stresses occur. (b) Conceptual illustration of expression distribution difference between distinct cell types and adjacent cell states. (c) Brief illustration of our work. scClassify2 adapts stable log-ratio values of expression data and incorporates prior gene co-expression knowledge via a dual-layer graph neural network (MPNN) to capture the expression topology of cells and then identify them accurately with a novel ordinal regression component. (d) scClassify2 could be applied to not only the traditional scRNA-seq data, but also sequencing data from subcellular spatial transcriptomics. (e) Brief illustration of the MPNN architecture adopted by scClassify2. Gene identities are encoded as nodes of the graph meanwhile the log-ratio of corresponding expression is encoded as edges. The model contains an encoder for topology capturing and a decoder for better learning.

Annotation between different single-cell expression data is subject to dataset or batch effect, where the distribution of gene expression varies across datasets due to technical and biological differences even after log transformation. This makes the generalizability of the single-cell classification model for cell annotation a persistent challenge. Selected classification methods such as CHETAH(21), scLearn(22), scMap(23) and scTyper(24) use correlation-based distance instead of Euclidean distance to match between query cells and training cells as correlation is unit independent. Other strategies involve using data integration strategies to find common embeddings(25, 26) between the query and reference. However, this entails additional computational expenses and needs to be repeated for each query dataset of interest. Some machine-learning-based approaches also tried to learn the generalised cell identity annotations from one experiment to another by semi-supervised training(15, 16). More recently, pre-trained models, especially large language models (LLMs) have emerged as the latest state-of-arts tools in multiple disciplines. LLMs are characterised by their ability to generalise. Methods have applied LLMs to single cell research including Geneformer(27) and scGPT(28). However, these models require computationally intensive training based on a massive number of databases and require additional adaptation to optimise for specific tasks.

Here, we proposed scClassify2, a novel and transferable framework for adjacent cell state identification across platforms. This is achieved with three key innovations. Firstly, the transferable component is achieved by adapting our recent cross-platform biomarker strategies (29) which identifies reference-free markers by examining log-ratio of expression values in multiple samples to capture consistent relationship between two genes. The relationship between genes is shown to be more stable across datasets compared to individual gene expression. Secondly, we use a dual layer architecture to enable joint learning from both the expression information and gene co-expression pattern derived from the log-ratio of genes. Specifically, to capture the gene co-expression pattern as well as the relationship between states, we utilised a type of graph neural network (GNN) named message passing neural network (MPNN) for the first time in single cell research. MPNN incorporates both node and edge information(30), unlike other types of GNN that focus on node features while ignoring edge information. Thirdly, to identify adjacent cell state transition, we adopt ordinal regression as the classifier of the network with a conditional training procedure. Using eight diverse datasets, we show our transferable dual layer model out-performed other cell annotation methods on transitional cell state identification tasks. Our framework is generalizable for both cell type and cell states annotation including spatially resolved data. A catalogue framework of cell state annotations containing pre-train models for various human tissues is made available via a user-friendly web server (https://shiny.maths.usyd.edu.au/scClassify_catalogue/) as a community resource.

## Material and methods

### Data collection

We included eight diverse scRNA-seq datasets (Table S1) which not only provide highly credible cell states of certain sequential biological processes but also have been cited by other cell trajectory research for performance evaluation. The datasets covered both human and mouse, different tissue types and cell types, diseased and healthy. For example, mouse gastrulation embryo dataset (31) was found at NCBI Gene Expression Omnibus (GEO: GSE171588). To control variables and focus on cell state classification, we only collected “Epiblast” cells (13377 in total). According to embryo development time, these cells are naturally divided into 6 states, i.e., E6.5 (17.01%), E6.75 (6.40%), E7.0 (37.89%), E7.25 (31.06%), E7.5 (6.91%) and E7.75 (1.73%). A detailed distribution of these cells could be found at Fig. 2a. We filtered cells which expressed less than 100 genes and genes which were expressed by less than 10 cells. No additional processing was conducted after collection and raw unnormalized expression count matrix was directly used by scClassify2.

**Fig. 2.**
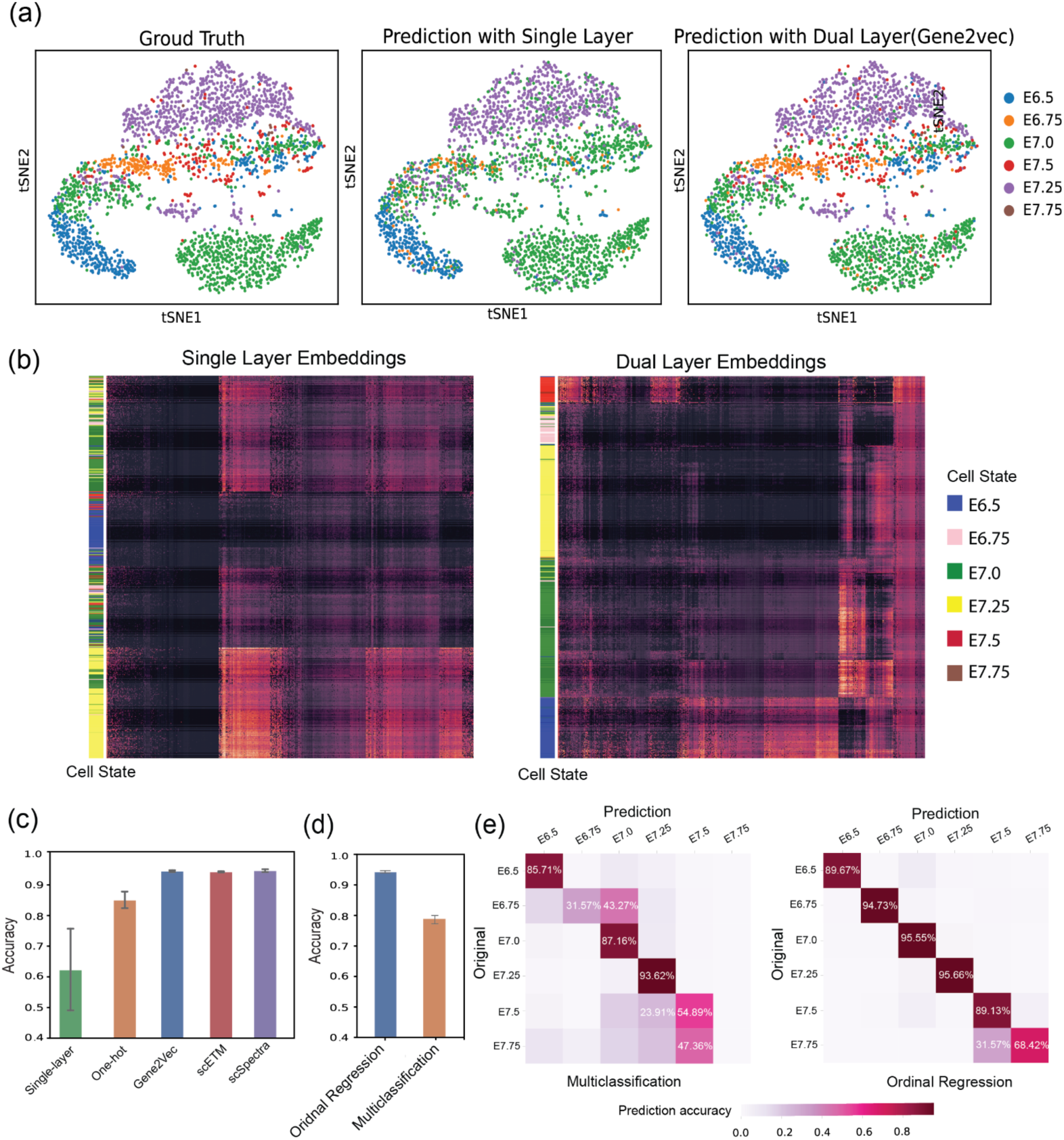
scClassify2 uses dual layer architecture based on message passing neural network. (a) tSNE plots of validation mouse gastrulation embryo dataset to show the prediction difference between single layer architecture and dual layer (with gene2vec). The left panel is annotated by ground truth. The middle and right panel are the prediction of single layer and dual layer architecture correspondingly. (b) Clustered cell embeddings captured by single layer (left panel) and dual layer (right panel) architecture correspondingly. In the heatmap, each row represents a cell, whose state is marked by the very left coloured stripes. As we can see, cell embeddings obtained from dual layer architecture show certain patterns which are highly correlated with corresponding cell state, while single layer ones do not. (c) Comparison among single layer using only gene expression data and dual layer integrating different sources of prior biological knowledge. (d) Prediction accuracy comparison between classifiers of general milti-classification and specifically designed ordinal regression across various cell states. (e) Confusion matrices of predictions from different classifiers show more details.

### Cell Graph construction

For each cell graph constructed by scClassify2, the input features include embedded nodes with gene representation and encoded edges with pairwise log-ratio value of gene expression counts. In particular, we use Gene2Vec(32), a distributed representation dictionary pretrained from transcriptome-wide gene co-expression data, to embed gene identities into node vectors with 200 dimensions.

As for edges, we firstly pick up around 600 highly variable genes and then transfer gene expression counts into pairwise log-ratio values. For a typical matrix ***X*** of gene expression counts with dimension ***n*** × ***m***, where ***n*** is the number of observations (cells), and ***m*** is the number of variables (sequenced genes). We define the “pairwise log-ratio values” for ***n*** cells as ***n*** square matrices all of size ***m*** × ***m***. More specifically, for each cell (assume its index is ***i***), the gene expression counts vector would be ***X_i_***, and the ***j***-th value ***X_i, j_*** is the expression counts of Gene ***j.*** For the ***m*** × ***m*** log-ratio matrix, its ***j***-th column would be enumerated by *logX_i, j_* − *logX_i.k_* for ***k*** in the range of ***1*** to ***m***, including ***j***. We add a Gaussian noise with mean of 0.5 to original expression counts to avoid computational failure caused by the data sparsity.

We used radial basis functions (RBFs) with Gaussian kernels to transform these single numerical ratio values into dense multidimensional vectors by mapping each input onto a high-dimensional space using radial symmetry.

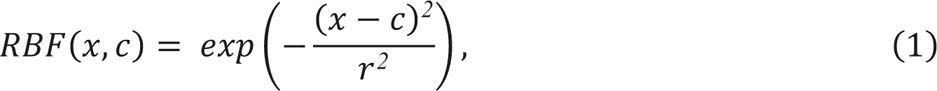

where *x*, *c* and *r* represent the log-ratio scalar needing to be vectorized, the centre point and the variance (spreadability) of the current Gaussian kernel respectively. We have calculated the distribution of these log-ration values and found most of them were within the range of -6 to 6. For efficient vectorization, we used 8 kernels whose centre points were unevenly spaced in that range according to our parameter searching process.

### MPNN

After comprehensively balancing the model performance and computation consumption like memory usage, we pruned the previous dense graph and only kept 16 outgoing edges for each node. More specifically, for each gene (node in the graph), we chose 8 other genes with largest log-ratio values and 8 with smallest values and kept corresponding edges. It is worth noting that log-ratio values between two paired genes are actually the opposites, i.e., if we mark the expression count of gene A and gene B as ***EA*** and ***EB***, then the log-ratio of A to B would be ***log(EA/EB)*** while B to A is ***log(EB/EA)***. To fully utilise the edge information, we added a perceptron layer for information propagation. This perceptron layer effectively integrates and reweights the corresponding neighbourhood of a certain node, serving as an attention mechanism and making the graph topology fit expression data better.

As shown in Fig1, the MPNN model of scClassify2 has an encoder-decoder architecture. The dual-layer encoder absorbs nodes and edges of the cell graph to gather messages from neighbourhoods and then alternatively updates nodes and edges by these messages passing along edges. More specifically, after aligning all input vectors, we first concatenate every two node vectors with the edge vector connecting them and calculate the message of this edge by a perceptron. Then we update node vectors using this message by a residual module with normalisation and dropout. With all nodes updated, we recalculate the message via another similar perceptron and then update edge vectors this time using new messages. The decoder is much simpler compared to the encoder. It takes nodes and edges from the encoder and computes messages along edges. The decoder only updates node features of the graph and tries to reconstruct the distributed representation of genes. Before we evaluated our method, we carried out hyper-parameter searching (Supplementary Fig, S2).

### Ordinal regression

Drawing on previous work(33), we construct ***n–1*** conditional training subsets ***S_1_***, . . . , ***S_j_***, . . . , ***S_(n−1)_*** for ***n*** sequential cell states. The first subset is the whole training set, and we conduct binary classification with regular binary cross-entropy loss to decide whether a cell belongs to the first state or following states. Meanwhile, the ***j^th^***subset only contains samples belonging to states following the (***j-1)^th^*** state and we also conduct similar binary classification with cross-entropy to decide whether a cell belongs to the ***j^th^***state or states following the ***j^th^*** state. After transferring the original large and complex multi-classification problem to a series of smaller but simpler binary classification problems, we obtain unconditional probability through the chain rule of conditional probability. More specifically, the predicted state index ***q*** of a cell would be

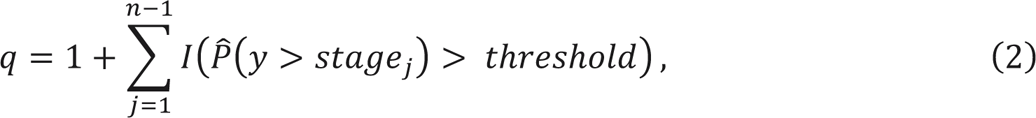

where ***I*** is the indicator function, *y* represents current cell state and *p̂*(*y* > *stage j*) means the estimated probability of the current cell belonging to states following the ***j^th^*** state, and the threshold is a tuneable predefined probability value. Then, the ordinal regression loss of scClassify2 would be:

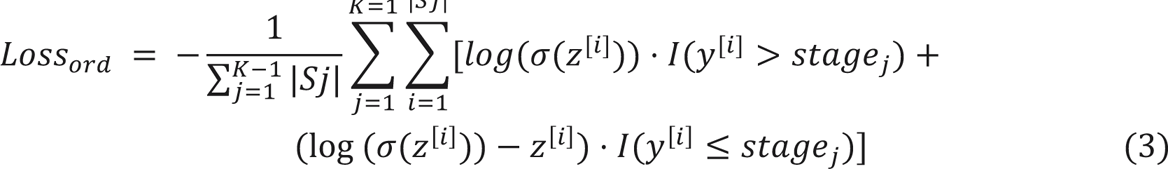

where *z*^[i]^ represents the predicted logits of the ***i^th^*** training example in ***S_j_***, *σ* means sigmoid function and ***S_j_*** means the ***j^th^***training subset with all cells belonging to states following the ***(j-1)^th^***state.

### Gene embedding reconstruction

Through passing messages along graph edges and self-updating, MPNN could concentrate locality into global gene expression topology, where subtle cell state information hides. To better assist in this topology capturing, we add gene representation reconstruction loss beside the cell state classification loss. The reconstruction loss measures overall distance between the distributed gene representation inputted to nodes with random masks (mask rate 15%) in the first layer of encoder and the corresponding recovered gene representation distribution outputted by the last layer of decoder. Since these two distributions might shift unevenly by computations and operations conducted within MPNN and thus do not share the same base anymore, we choose an approximating Wasserstein distance(34–37) as our loss instead of directly using Kullback–Leibler divergence(38). The reconstruction loss would be:

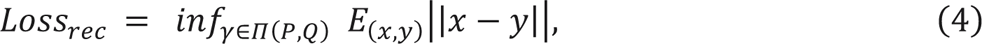

where *γ* is the joint distribution of original gene representation ***P*** and reconstructed gene representation ***Q***, *Π* is the set of all possible *γ*, ***inf*** means the infimum of the expectation.

### Loss function

As described above, the loss function of scClassify2 has two main parts: an ordinal regression part to directly classify cell states according to the gene expression pattern it captures, and a gene embedding reconstruction part to implicitly guide the topology capture of MPNN and thus benefit the previous one. We use two empirical coefficients to combine them:

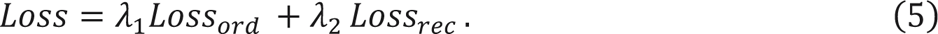

### Dual-layer implementation

We compared the single-layer network which only used expression counts with the dual-layer architecture which integrates estimated gene-gene association information with gene expression. As mentioned above, an observation or a cell is represented by a graph, where the nodes represent genes and the edges represent log-ratio values of two connected genes’ expression counts. For implementation convenience, we kept the main framework and designed the comparison of node fillers in order from simplicity (no information) to complexity (more information), where each node could be

a. set to all zeros - representing no information in nodes and the network only used expression counts.
b. one-hot vectors - representing only unrelated gene information in nodes.
c. gene embeddings from Gene2vec(32) - representing gene-gene association information estimated by a machine learning approach from large-scale gene-gene co-expression data.
d. gene embeddings from scETM(39) and scSpectra(40) - representing gene-gene association information estimated by deep learning approaches from transcriptome-wide gene-gene co-expression data.

### Evaluation framework

We mainly used accuracy and confusion matrix for our performance evaluation. A confusion matrix is a table to visualise the detailed cell state identification performance. It presents a summary of the percentages of correct and incorrect predictions made by the model, organised by the ground truth and predicted labels. We executed strict 5-fold cross validation on a collection of eight datasets (Supplementary Table S1) covering both human and mouse, different tissue types and cell types, diseased and healthy. Before the systemic evaluation, we checked the stability of our method (Supplementary Fig.S3).

To evaluate and compare the performance of scClassify2, we obtained four other publicly available methods by their default settings. These packages were installed either through their official Bioconductor website or directly from their GitHub page before December of 2023. Among them, scClassify is our previous version, which is based on traditional ensembled k-neighbours network (KNN) and cell type hierarchies. sigGCN is a multimodal end-to-end graph convolutional network combined with gene interaction information. scGCN is another graph convolution network which achieves effective knowledge transfer across disparate datasets. scGPT is a pre-trained foundation large language model with fine tuning pipeline for single-cell multi-omics.

### Subcellular spatial transcriptomics (SST)

We used the Xenium Breast Cancer dataset downloaded from https://www.10xgenomics.com/products/xenium-in-situ/preview-dataset-human-breast to validate scClassify2’s capability on SST data. This process needed to combine Bidcell, a cell segmentation tool developed by our group. We used its default config (details could be found at: https://github.com/SydneyBioX/BIDCell) to get the cell segmentations and applied scClassify2 on segmented expression files.

### Web server

For the front-end, the interface of our web server, scClassify-catalogue (https://shiny.maths.usyd.edu.au/scClassify_catalogue/) is based on Shiny. Shiny is an R package facilitating the creation of interactive web applications, enabling seamless integration of data analysis and visualisation for end users. Meanwhile, the back-end functionality was implemented through Python, as described in our Github page (https://github.com/Wenze18/scClassify2).

## Results

### scClassify2 learns expression topology by integrating prior biological information through message passing network

We introduce scClassify2, which provides a generalizable and transferable cell annotation framework for sequential cell state identification across platforms. With the increased number of perturbation studies, this new extension of scClassify is specifically designed to recognise the inherent order of cell state. In our previous study, our scClassify captures the hierarchical nature of the cell types through a cell type tree(41). To further increase our capacity to effectively distinguish subtle cell type or cell state differences, we used a dual-layer design with a type of graphical neural network called message passing neural network (MPNN). We chose MPNN as the backbone due to its ability to capture both node and edge information as well as its flexibility and scalability (42–45). Based on MPNN, we encode each cell as a graph and employ a dual-layer deep learning approach, as illustrated in Fig1. The dual-layer design allows the integration of two levels of information in the network, i) log-ratio of pairwise gene expression counts modelled as edge and ii) biological knowledge derived from gene co-expression modelled as node. This allows information to propagate among genes across connecting edges and captures subtle gene expression topology of different cell states including gene co-expression. We used the ratio of gene expression as it is shown to be relatively more stable across datasets compared to individual gene expression (29).

To assess the benefit of additional gene information in enhancing cell state annotation, we compared the single-layer mode with the dual-layer mode, where the former relies solely on expression data. As illustrated in Fig. 2a, the integration of biological information into gene representations via the dual-layer architecture yields higher accuracy for sequential cell state identification compared to no information (e.g., the accuracy of using one-hot vectors versus zero vectors is 0.63 versus 0.86). Moreover, we checked the cell embeddings captured by MPNN correspondingly (Fig. 2b) and found dual-layer information could help the neural network recognize and construct specific embedding patterns of subtly different cell states much better.

We further investigated model performance using various implementations to capture the gene co-expression pattern. We found the performance of cell state identification could be further enhanced by utilising distributed gene representations as node embeddings. For instance, as depicted in Fig. 2c, the accuracy of cell state identification increases from 0.86 when using one-hot vectors to 0.95 when employing vectors from Gene2vec. Such distributed gene representations, derived from embedding methods like Gene2vec, scEMT, and scSpectra, estimate gene regulation networks by capturing gene co-expression patterns from large-scale transcriptome-wide data. Gene2vec, for example, derives the pattern using nearly 1000 datasets from GEO. Based on the performance of these embedding methods, we implemented Gene2Vec in scClassify2 to embed each gene into a node vector with 200 dimensions.

### scClassify2 uses ordinal regression to effectively identify adjacent cell states

Given the many biological processes that involve transitional cell states, it is important to capture the sequential nature between transitional cell states, scClassify2 introduced an ordinal regression layer in the model and a novel training procedure based on the conditional probability distribution of adjacent cell states.

To demonstrate its effectiveness, we compared scClassify2 with a conventional multi-classification layer on a mouse gastrulation embryonic development cell state dataset. Fig. 2d highlights the improved performance of the ordinal regression with a cell state prediction accuracy of 0.93 compared to 0.82 of the conventional multi-classification. Notably, the multi-classification model had trouble identifying E6.75. It only correctly identified ∼30% of E6.75 and incorrectly predicted over 40% as E7.0. In contrast, the cell classification model based on ordinal regression in scClassify2 can correctly identify almost 95% of the E6.75 as E6.75 cells (Fig. 2e). Given the subtle differences in gene expression profiles along adjacent cell states, the use of ordinal regression and the conditional probability distribution between states underscores the benefits of learning the underlying relationship between the cell states.

### scClassify2 outperforms other advanced approaches for sequential cell state identification

We compared the performance of scClassify2 with our previous version, as well as three other state-of-the-art classification methods sigGCN, scGCN, and scGPT, across eight datasets (Table S1) on sequential cell state identification task (Fig.3). scClassify2 represents a significant improvement over our previous work, scClassify, evident across all eight datasets. For example, on dataset 8, scClassify2 achieves a prediction accuracy of 80.82 ± 2.14% compared to 66.44 ± 4.19% for scClassify. When compared with other state-of-the-art graph-neural-network-based methods such as sigGCN and scGCN, scClassify2 demonstrates consistent performance advantages across all datasets. For example, as shown in Fig.3, scClassify2 achieves an accuracy of 88.91 ± 1.73%, outperforming sigGCN and scGCN with 79.32 ± 1.76% and 77.50 ± 7.17% respectively on dataset 3. Notably, scClassify2 slightly outperforms scGPT, the latest cell annotation method using generative artificial intelligence, on most test datasets. For example, on dataset 1, scClassify2 achieves an accuracy of of 94.50 ± 0.39% compared to 93.02 ± 0.42% for scGPT), outperforming the model (scGPT) that is pre-trained on thousands of information, a repository of over 33 million cells, providing it with a vast amount of prior knowledge. In contrast, scClassify2 is trained on specific tissue dataset, which provides it with the flexibility of tailoring the reference dataset than generalised models.

**Fig. 3.**
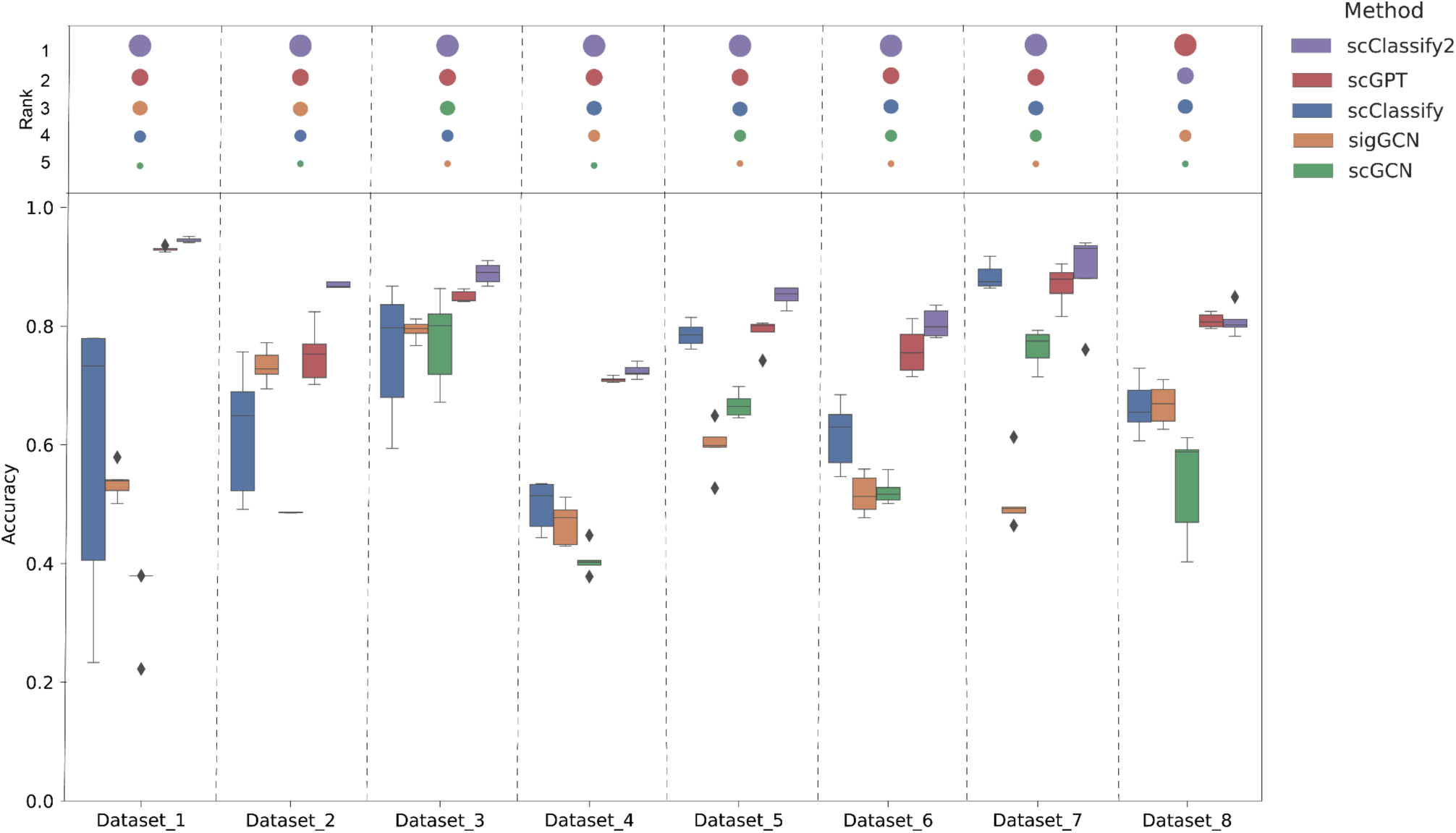
Performance comparison for eight different datasets. Comparison of model performance for 5 advanced approaches on 8 different sequential cell state dataset. Relative ranks of these methods on each dataset are presented at the top.

### Broad applicability of scClassify2 including subcellular spatial transcriptomics data

To assess the applicability of scClassify2 on other biotechnology platforms, we tested it on a breast cancer dataset generated by a subcellular spatial transcriptomics (SST) platform, Xenium. SST is the latest spatial technology in the new era of spatial omics sequencing. We firstly applied BIDCell, our recently published state-of-the-art segmentation algorithm, to delineate individual cells. As illustrated in Fig. 4a-d, scClassify2 worked well even for scenarios without sequential relationships among cell types and achieved an identification precision of around 0.92 when compared to scClassify. Furthermore, we found the prediction accuracy of scClassify2 is relatively consistent across spatial regions of the breast cancer slide (Fig.4e-g), while slightly fluctuating among different cell types (Fig. 4h). Our findings prove scClassify2 as a cell type identification method with broad applicability that can be seamlessly applied to SST data.

**Fig. 4.**
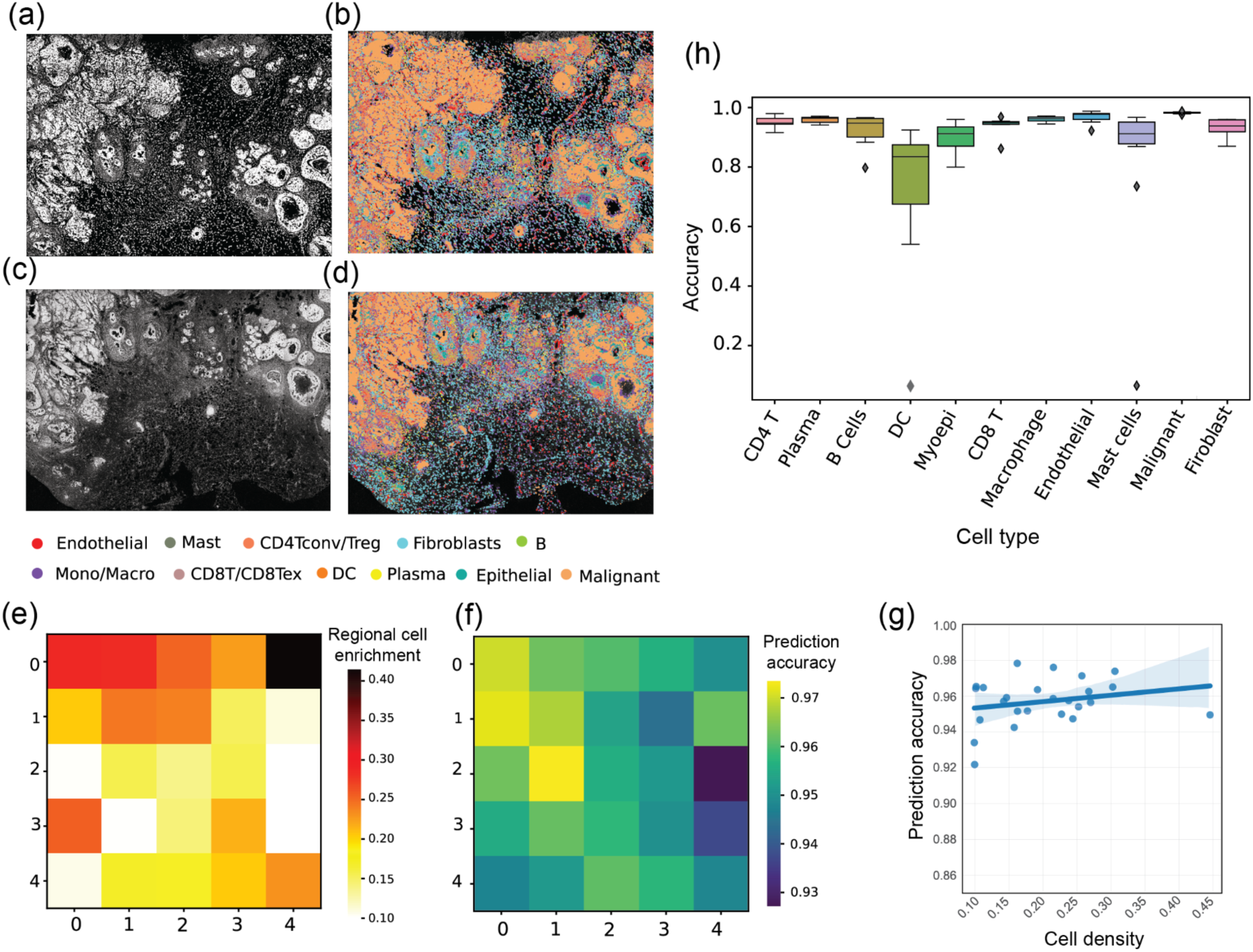
Applicability of scClassify2 for SST data. scClassify2 could be directly applied on subcellular spatial transcriptomics (SST) data. (a)-(d) The raw image and cell types identified by scClassify2 of human breast tissues, where (a) and (b) is replicate 1 while (c) and (d) stands for replica 2. (e)-(g) We equally divided the vision into 25 regions to observe the annotation results of scClassify2. (e) Regional cell enrichment (how many cells in one specific region compared with the whole vision). (f) scClassify2’s prediction accuracy for each region. (g) The relationship between regional cell enrichment (x-axis) and scClassify2’s prediction accuracy (y-axis). (h) We also checked scClassify2’s precision for each cell type when applied it on SST data.

### scClassify2 provides catalogue for convenient single cell annotation

To provide a resource for the scientific community, we developed a user-friendly web server called scClassify-catalogue, offering a comprehensive catalogue of scClassify2 models trained from datasets covering almost 1, 000 cell types and 30 tissue types (Fig. 5a). Traditional web portals often present multiple non-integrated references for the same tissue and place overhead on the users with the task of experimenting with multiple references. In contrast, scClassify-catalogue directly delivers annotation results for the selected tissue within a relatively short time. The input of scClassify-catalogue is flexible and allows either the direct output from 10x cell ranger software, or a csv file or h5ad file containing the gene expression matrix (Fig. 5b). Once the job is submitted, a job ID will be made available (Fig. 5c-d). A file containing the predicted outcome as well as an HTML file will be emailed to the email address entered by the user once the job finishes. The HTML file visualises the predicted cell type such that the user can easily check the prediction result (Fig. 5e).

**Fig. 5.**
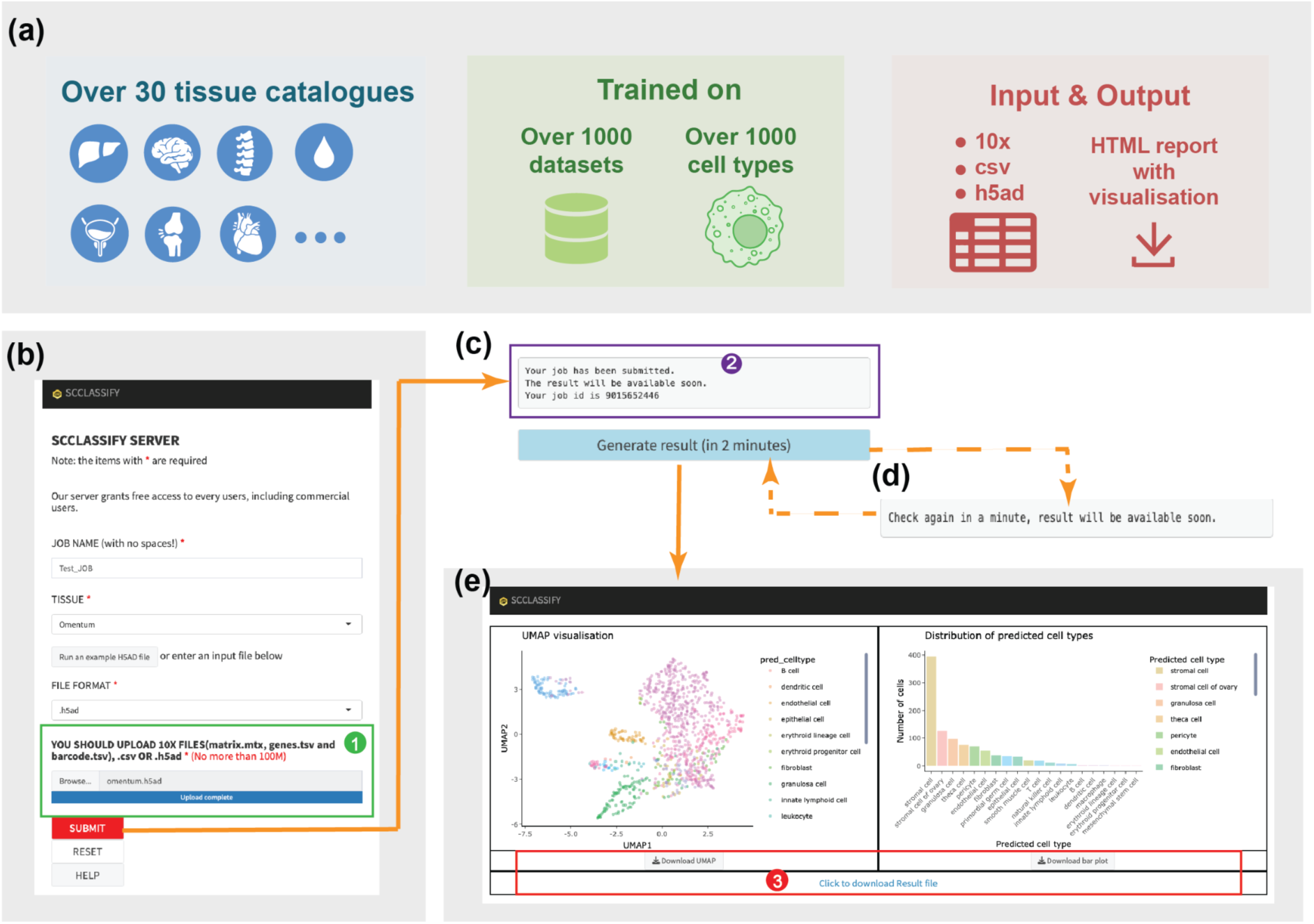
Overview of scClassify-catalogue - the web server. (a) A brief overview of our web server, scClassify-catalogue. (b) Homepage of scClassify-catalogue. The window for uploading expression profiles of query data is marked by a green rectangular (Box 1). After filling all necessary information in, click the “SUBMIT” button. (c) After submission, the waiting panel would pop up. The prompt window of the waiting panel is marked by a purple rectangular (Box 2). Job ID in this window is important for us to trace the backend activity of the user’s submission. Please wait enough time (usually less than 2 minutes) before clicking the “Generate result” button. (d) If the job has not been finished, another waiting panel would pop up. (e) The result page of the finished job. A UMAP of input data with annotations and a brief statistical bar plot of analysis results is available on this page. Downloadable and editable result table and analysis plot mentioned above could be reviewed by users Box 3).

## Discussion

In this study, we introduce scClassify2, a cell state identification method based on log-ratio values of gene expression, a message passing framework with dual-layer architecture and ordinal regression. scClassify2 effectively distinguishes adjacent cell states with similar gene expression profiles. Our comprehensive evaluation demonstrates the superior performance of scClassify2 compared to other state-of-the-art methods including scGPT, sigGCN and scGCN. Moreover, scClassify2 can be applied for cell annotation tasks on SST data. To enhance its accessibility for researchers, we have developed a user-friendly web server equipped with pre-trained models for over 30 tissues.

The integration of dual-layer learning which combines expression data with prior biological knowledge represents an innovative strategy to incorporate prior knowledge. As intended, relevant additional information leads to improved prediction performance. Using distributed gene representations such as Gene2vec, scETM and scSpectra as node embedding lead to further enhancement in performance. These representations estimate gene regulation networks by capturing gene co-expression patterns from transcriptome-wide data and play a critical role in enabling the network to discern subtle high-dimensional expression pattern disparities among neighbouring cell states. Interestingly, the results suggest that MPNN is quite an effective learning model as the simpler gene embeddings from Gene2vec is sufficient to achieve best performance compared to more comprehensive and detailed association information from scETM and scSpectra.

The distinct efficacy of MPNN in comparison to other graph neural network architectures (Fig.S1) lies in its unique message passing mechanism, enabling nodes within the graph to exchange information along edges and influence neighbouring nodes. In contrast, Graph Convolutional Network (GCN) rely solely on aggregated signals within each node for self-update, potentially limiting their effectiveness. Despite attempts to augment GCN with attention mechanisms, such as Graph Attention Network (GAT) and graph transformer, they still fall short of MPNNs due to their inherent lack of message passing and propagation. This crucial process is indispensable for capturing subtle gene expression differences among adjacent cell states. With genes as nodes and pairwise log-ratio values of gene expression counts as edges, we speculate the graph itself mimics the whole signal pathway network within a cell, where messages of different layers computed by MPNN establish bridges facilitating gene communication akin to regulatory molecules in biological systems.

Since gene expression profile exhibits only subtle differences along adjacent cell states (Fig.1b), the improvement from multi-classification to ordinal regression suggests establishing a comprehensive consistency of gene expression distributions from different cell states rather than separately assigning them could leverage the nature of cell state evolution by using conditional probability among them and thus results in better identification.

We have shown that besides scRNA-seq, scClassify2 is directly applicable on SST. This offers the potential to continuously generate additional pre-trained models within the catalogue to accommodate other forms of spatial datasets including spatial proteomics, imaging mass cytometry (IMC) and spatial transcriptomics data. Spatial proteomics and IMC, different to transcriptomics data, sequence selective panels of genes containing only a few hundred genes. This requires some special adaptation strategy of the pre-trained models such as subsetting the models to the gene panels.

In summary, we present scClassify2, a single-cell annotation tool with an emphasis to identify sequential cell states in an ordinal manner that is applicable to both scRNA-seq and SST data. As a resource for the community, we have constructed a dedicated web server with pre-trained references available at (https://shiny.maths.usyd.edu.au/scClassify_catalogue/). Our work contributes to the community by alleviating the laborious process of selecting appropriate reference data and the requirement of computational infrastructure such as GPUs.

## Ethics statement

Not applicable.

## Data Availability

Details of each dataset used in this study is listed in Supplementary Table 1.

## Acknowledgments

The authors thank all their colleagues, particularly at The University of Sydney, Sydney Precision Data Science and Charles Perkins Centre for their support and intellectual engagement. Special thanks to Zhenghao (Chris) Chen for his contribution in our weekly discussion.

## Author Contribution

J.Y. conceived and funded the study. J.Y., W.D., Y.C. and X.F. completed the design of study. W.D and Y.C. completed the evaluation and analysis. The implementation of the package was done by W.D. The construction of the web server was done by M.T. All authors wrote, reviewed, and approved the manuscript.

## Supplementary Figures

**Supplementary Fig. S1.**
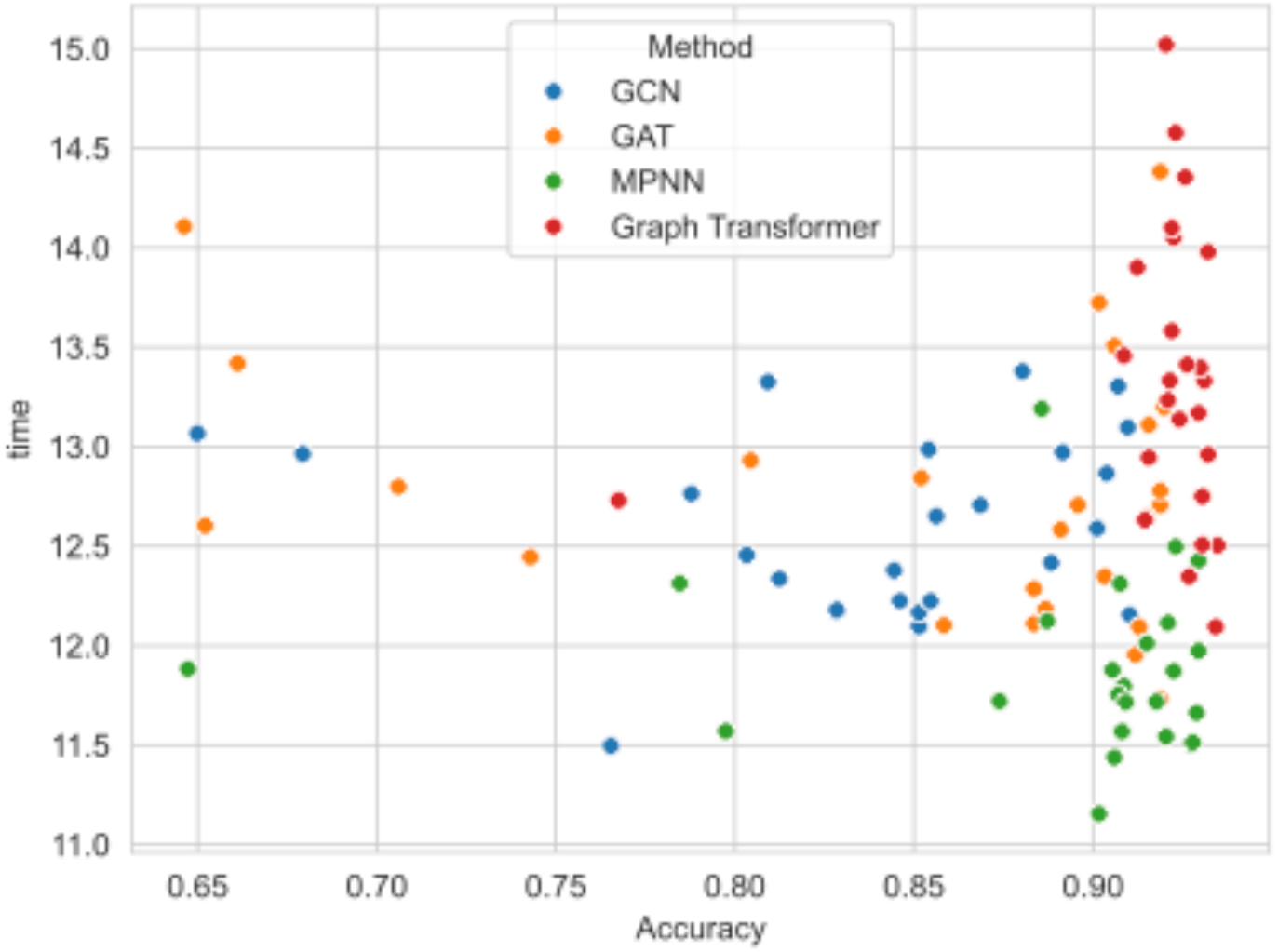
The comparison among several graph neural network architectures. The overall prediction accuracy of MPNN and graph transformer is approximately equal, outperforming GCN and GAT. However, the graph transformer consumes significantly more training time compared with MPNN.

**Supplementary Fig. S2.**
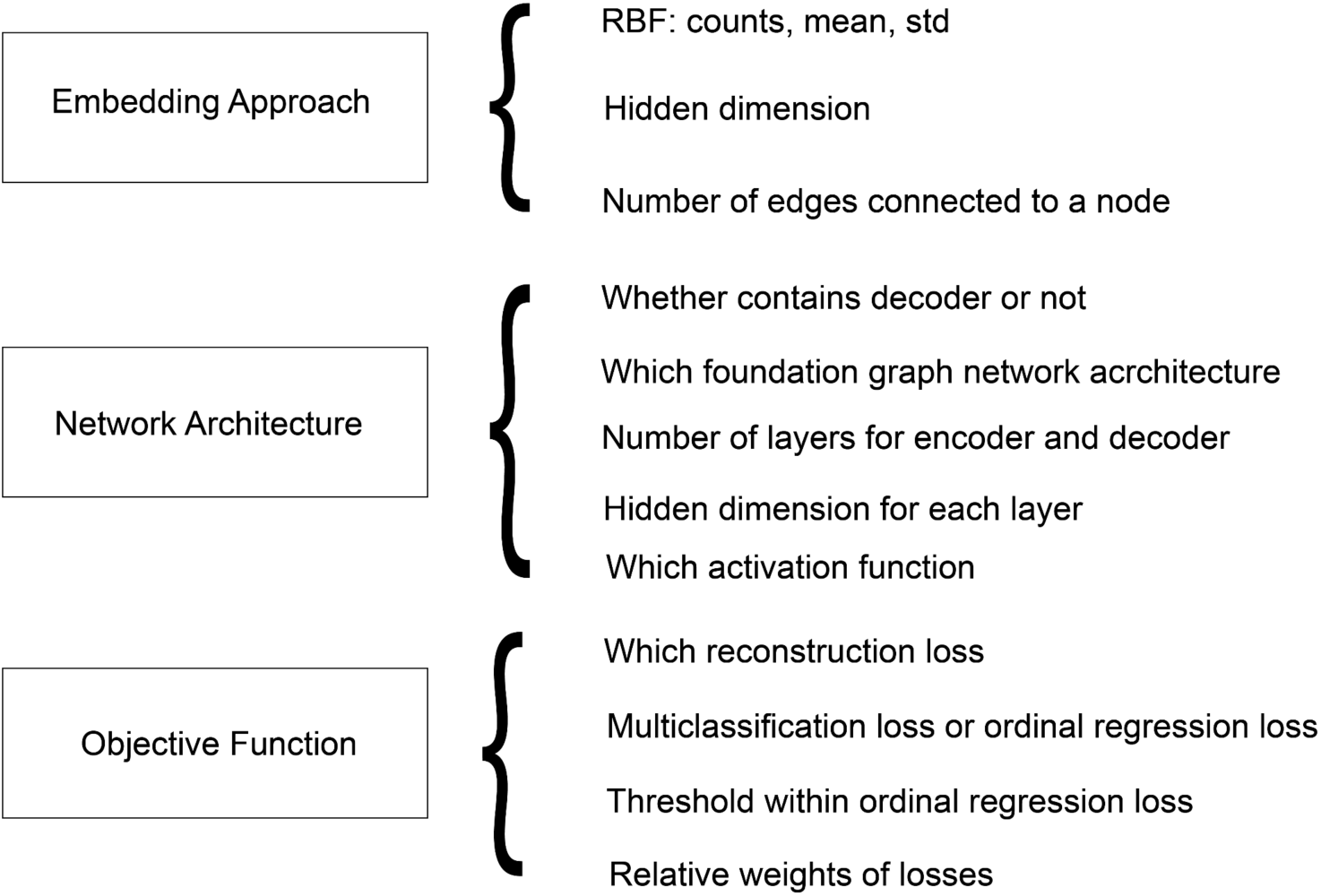
The comprehensive hyperparameter search was conducted for scClassify2. Network hyperparameters of 12 aspects from 3 domains were involved to optimise its performance and robustness for real-world applications.

**Supplementary Fig. S3.**
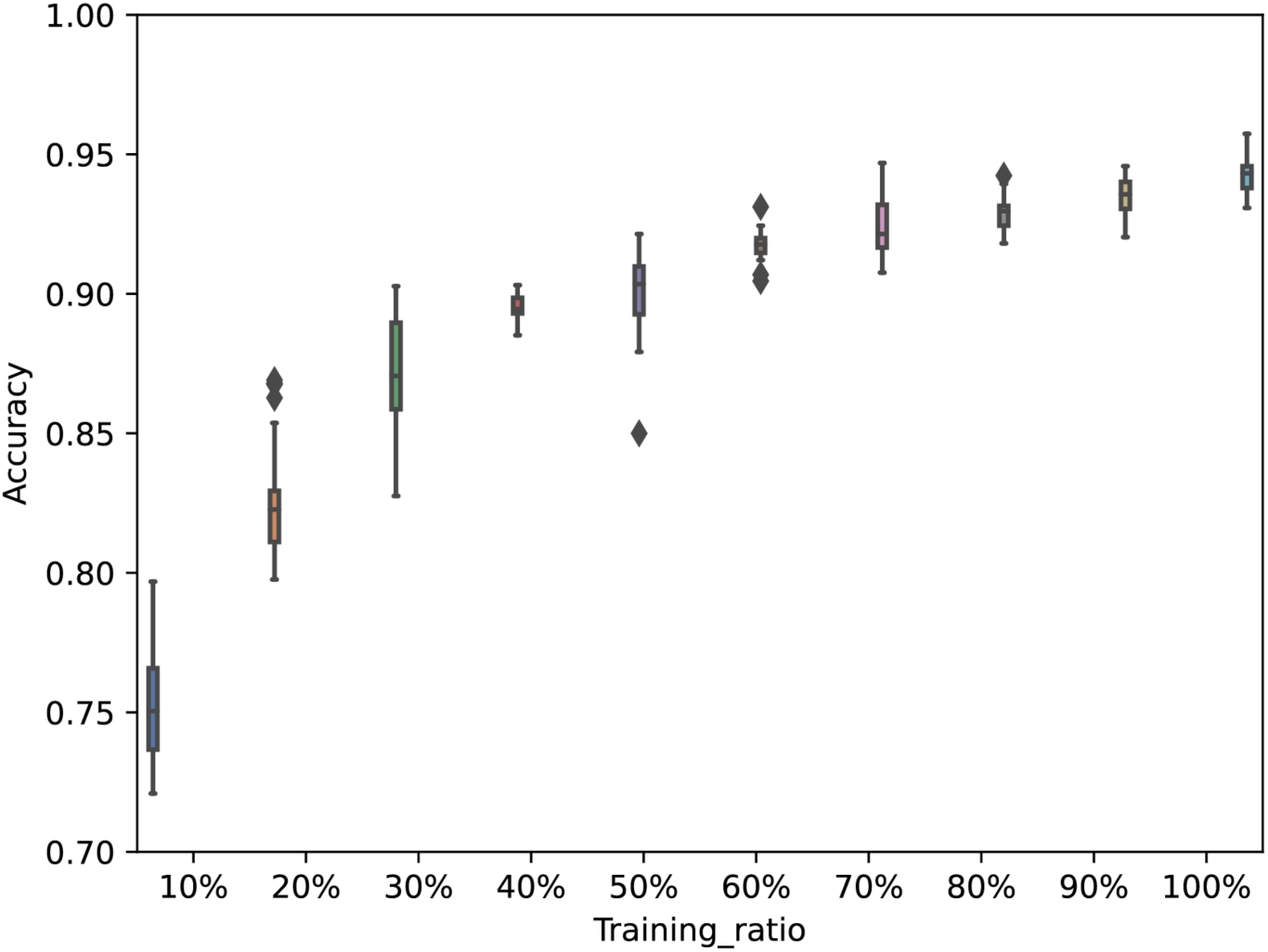
The stability of model performance with respect to changes in the size of the training set.

## Supplementary Table

**Supplementary Table S1.**
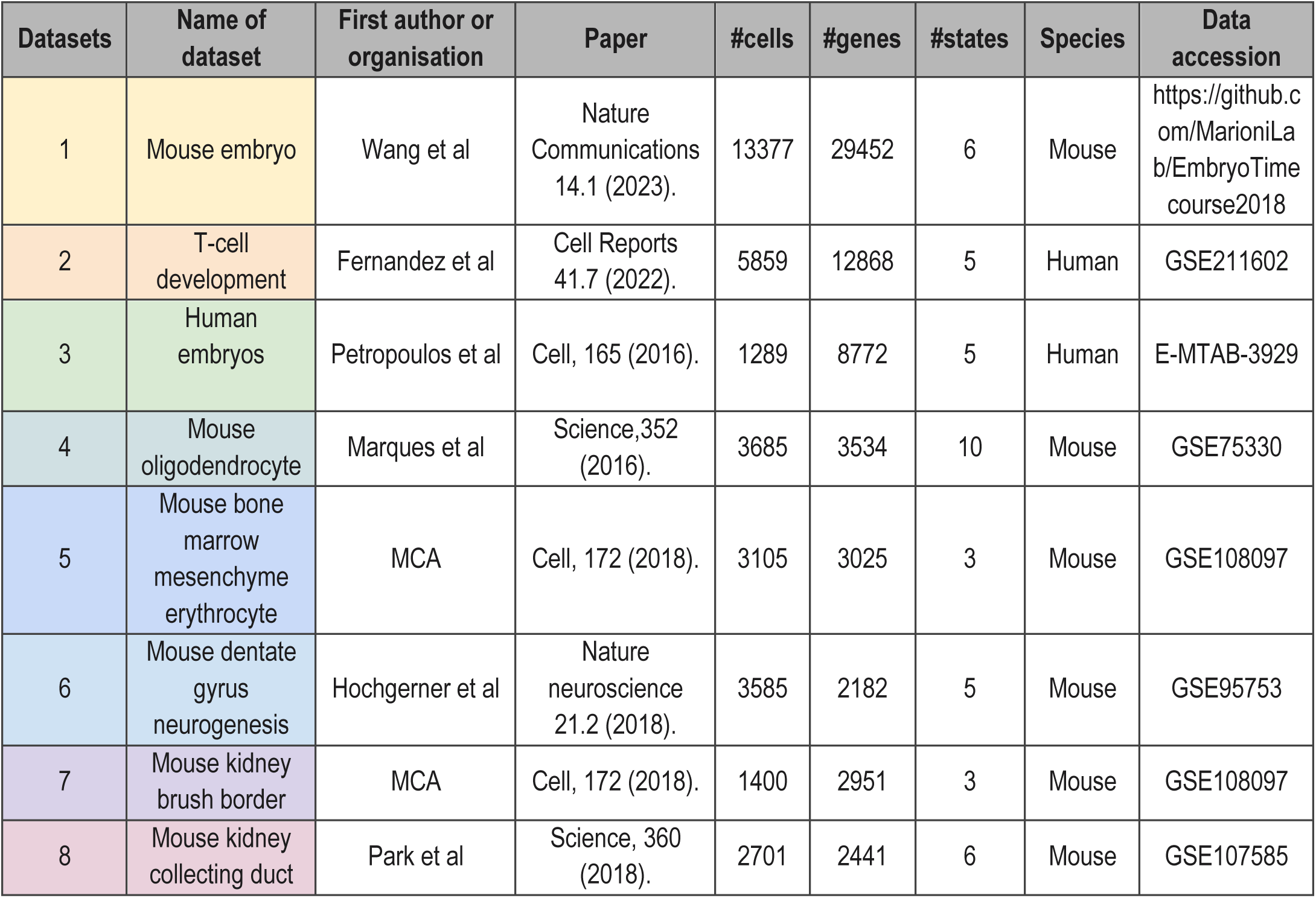
Details of eight scRNA-seq datasets of sequential cell state transformation. The columns #cells, #genes, #states denote the number of cells, the number of genes and the number of cell states respectively.

| Datasets | Name of dataset | First author or organisation | Paper | #cells | #genes | #states | Species | Data accession |
| --- | --- | --- | --- | --- | --- | --- | --- | --- |
| 1 | Mouse embryo | Wang et al | Nature Communications 14.1 (2023). | 13377 | 29452 | 6 | Mouse | https://github.com/MarioniLab/EmbryoTimecourse2018 |
| 2 | T-cell development | Fernandez et al | Cell Reports 41.7 (2022). | 5859 | 12868 | 5 | Human | GSE211602 |
| 3 | Human embryos | Petropoulos et al | Cell, 165 (2016). | 1289 | 8772 | 5 | Human | E-MTAB-3929 |
| 4 | Mouse oligodendrocyte | Marques et al | Science, 352 (2016). | 3685 | 3534 | 10 | Mouse | GSE75330 |
| 5 | Mouse bone marrow mesenchyme erythrocyte | MCA | Cell, 172 (2018). | 3105 | 3025 | 3 | Mouse | GSE108097 |
| 6 | Mouse dentate gyrus neurogenesis | Hochgerner et al | Nature neuroscience 21.2 (2018). | 3585 | 2182 | 5 | Mouse | GSE95753 |
| 7 | Mouse kidney brush border | MCA | Cell, 172 (2018). | 1400 | 2951 | 3 | Mouse | GSE108097 |
| 8 | Mouse kidney collecting duct | Park et al | Science, 360 (2018). | 2701 | 2441 | 6 | Mouse | GSE107585 |

